# Rapid Establishment of Tracheal Stenosis in Pigs Using Endotracheal Tube Cuff Overpressure and Electrocautery

**DOI:** 10.1101/2020.07.16.199679

**Authors:** Jin Hyoung Kim, Jong Joon Ahn, Yangjin Jegal, Kwang Won Seo, Seung Won Ra, Byung Ju Kang, Soohyun Bae, Soon Eun Park, Moon Sik Jung, Ju Ik Park, Hee Jeong Cha, Yongjik Lee, Taehoon Lee

**Affiliations:** Division of Respiratory and Critical Care Medicine, Department of Internal Medicine, Ulsan University Hospital, University of Ulsan College of Medicine, Ulsan, Korea; Department of Anesthesiology and Pain Medicine, Ulsan University Hospital, University of Ulsan College of Medicine, Ulsan, Korea; Department of Pathology, Ulsan University Hospital, University of Ulsan College of Medicine, Ulsan, Korea; Department of Thoracic and Cardiovascular Surgery, Ulsan University Hospital, University of Ulsan College of Medicine, Ulsan, Korea

**Keywords:** pig tracheal stenosis model, cuff overpressure, electrocautery

## Abstract

**Background:** Central airway obstruction can be caused by cancer, tracheal intubation, or tuberculosis, among others. If surgery is contraindicated, bronchoscopic therapy may be performed. Bronchoscopic treatment for airway obstruction is continuously evolving. In particular, attempts to overcome the current shortcomings of airway stents (stent migration, mucostasis, and granulation tissue formation) are currently ongoing. To apply a new airway stent to humans, preclinical studies in an appropriate animal model is needed. Canine and porcine tracheas have been used as animal airway stenosis models. However, existing models take a long time to develop (3–8 weeks) and have a disadvantage that the mechanism of stenosis is different from that in humans.

**Purpose:** To establish a new and fast tracheal stenosis model in pigs using a combination of cuff overpressure intubation and electrocautery.

**Methods:** Fourteen pigs were divided into three groups: tracheal cautery (TC) group (n = 3), cuff overpressure intubation (COI) group (n = 3), and COI-TC combination group (n = 8). Cuff overpressure (200/400/500 mmHg) was applied using a 9-mm internal diameter endotracheal tube. Tracheal cautery (40/60 watts) was performed using a rigid bronchoscopic electrocoagulator. After intervention, the pigs were observed for 3 weeks and bronchoscopy was performed every 7 days. When the cross-sectional area decreased by > 50%, it was judged that tracheal stenosis was established.

**Results:** The time for tracheal stenosis was 14 days in the TC group and 7 days in the COI-TC combination group. In the COI group, no stenosis occurred. In the COI-TC group, electrocautery (40 watts) immediately after intubation for > 1 hour with a cuff pressure of 200 mmHg or more resulted in sufficient tracheal stenosis within 7 days. Moreover, the degree of tracheal stenosis increased in proportion to the cuff pressure and tracheal intubation time.

**Conclusions:** The combined use of cuff overpressure and electrocautery helped to establish tracheal stenosis in pigs rapidly. This animal model was technically simple and reproducible, and used a mechanism similar to that in human tracheal stenosis. It is expected to help develop new treatments for airway stenosis

## INTRODUCTION

Central airway obstruction (CAO) is broadly defined as obstruction of the trachea, either of the main stem bronchus and/or the bronchus intermedius (1). It might occur secondary to tumors, tracheal intubation, or infection. CAO can frequently cause dyspnea, asphyxiation, and even death. In cases of benign CAO, surgery is the treatment of choice. However, in cases with contraindication to surgery, bronchoscopic intervention is a potentially useful treatment option. Similarly, in malignant CAO, surgery can be performed if operable, but bronchoscopic intervention is performed in cases of inoperable advanced cancer or terminal cancer.

A variety of endoscopic procedures are available to alleviate CAO. Balloon dilation, airway stent, mass excision (debulking), and tumor ablation (laser therapy, cryotherapy, argon plasma coagulation, electrocautery, and photodynamic therapy) have been used (2). However, the management of CAO remains difficult. Besides other techniques, stent-related technologies continue to evolve to overcome the current shortcomings of stents (migration, mucostasis, and granulation tissue formation). Recently developed stents or currently developing stents include drug-eluting stents, stents with a new design, stent with a hydrophilic coating of the inner surface, radiopaque silicone stents, bioabsorbable stents, and 3D-engineered/printed stents (3-6).

To investigate and develop a new effective treatment for CAO, it is necessary to establish an experimental animal model. Several animal (dog, pig, and rabbit) models of tracheal stenosis have been developed through various methods such as chemical caustic agents, laser, and electrocautery (7-10). However, these methods are sophisticated and require a relatively longer time (3–8 weeks) to induce stenosis. Moreover, these models are inconsistent with the actual mechanism of human tracheal stenosis. Prolonged intubation and tracheostomy are the most common causes of human benign tracheal stenosis. The actual mechanisms underlying this stenosis include a combination of (1) excessive cuff pressure, (2) over-sized intubation tube, and (3) tracheal injury. Local ischemia caused by either excessive cuff pressure or an over-sized intubation tube and tracheal injury (trauma during intubation, or tracheostomy related injury) can result in regional inflammation and subsequent stenosis during healing (11-13).

Recently, Su et al. reported that prolonged intubation (24 hours) with cuff-overpressure and an over-sized tube could induce tracheal stenosis in dogs in a relatively shorter time (2 weeks)(14). This study inspired our research. We hypothesized that if three mechanisms (excessive cuff pressure, over-sized intubation tube, and tracheal injury) are applied at the same time, it would be possible to establish an animal tracheal stenosis model more quickly.

The purpose of the present study was to establish a new and rapid animal model of tracheal stenosis in pigs through the combined use of prolonged over-sized intubation with cuff-overpressure and tracheal injury (tracheal cautery), and investigate the difference in the incidence and speed of tracheal stenosis development according to the degree of cuff overpressure, intubation duration, and tracheal injury (tracheal cautery).

## METHODS

### Animals

Twelve-week-old female farm pigs (weight, 40–45 kg; tracheal size, 16–20 mm) were used because their tracheal size is similar to that of humans. The animal experiments were conducted at the preclinical trial center of Pusan National University Yangsan Hospital. The experimental animals were handled according to well-established bioethical guidelines (Guiding Principles in the Care and Use of Animals, DHEW Publication, NIH), following a protocol approved by the Institutional Animal Care and Use Committee at Pusan National University Yangsan Hospital (Approval Number: PNUYH-2017-043).

### General anesthesia

Intramuscular alfaxan (5 mg/kg) and inhalational 3% isoflurane were used for the induction and maintenance of general anesthesia, respectively.

### Tracheal cautery (TC)

Tracheal cautery (40–60 watts) was performed using a coagulation suction tube (10390 BN, Karl Storz, Germany) via a rigid bronchoscope (size 8.5, 10318 BP, Karl Storz, Germany) (Figure 1). The cautery was performed at eight sites (including the membranous portion) of the tracheal segment usually 5 cm below the vocal cords at a length of 1 cm parallel to the trachea. Cautery was performed after (prolonged) cuff overpressure intubation, at the position where the cuff was located.

**Figure 1.**
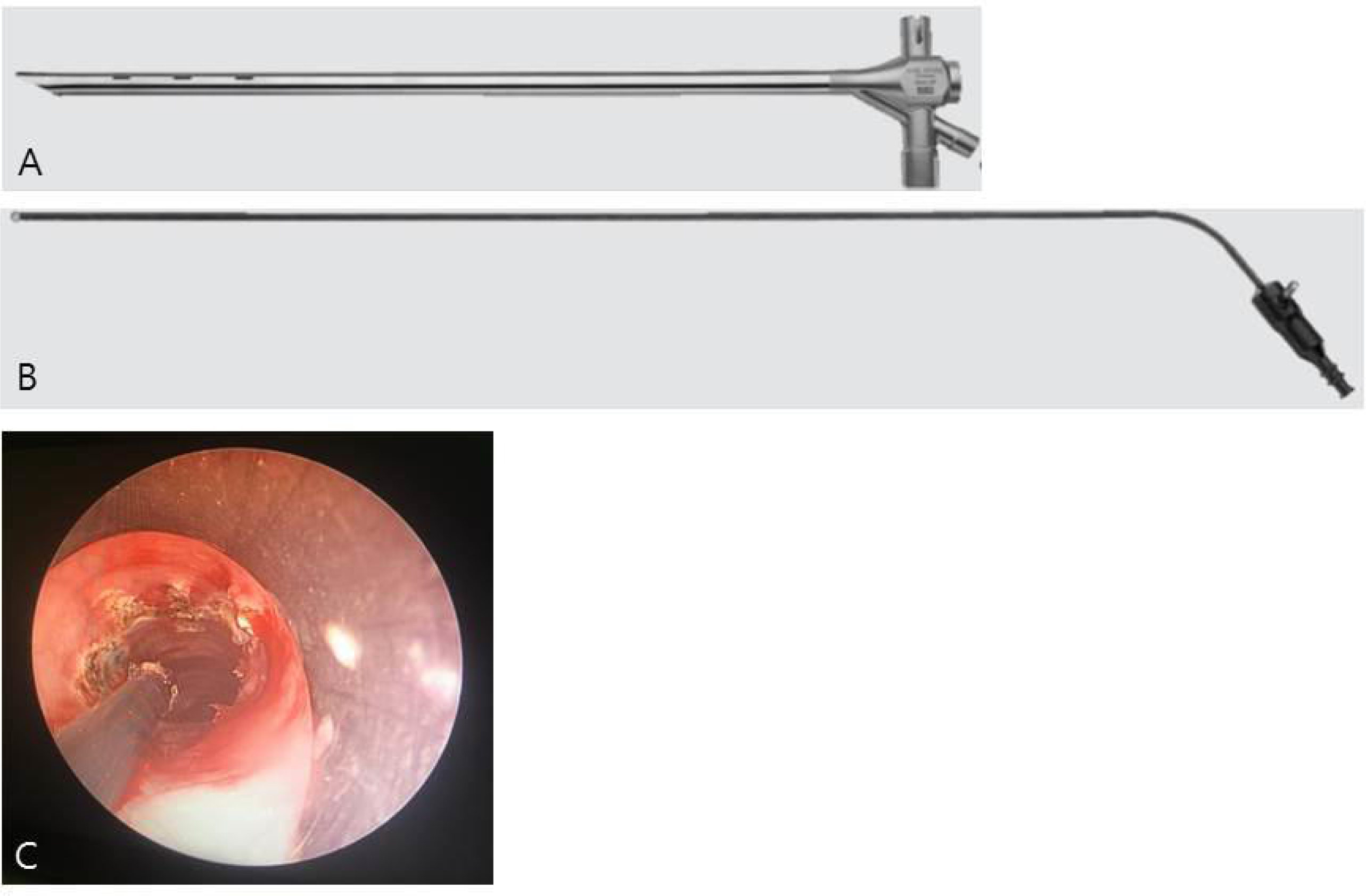
The equipment used for tracheal cautery. (A) Rigid bronchoscope (size 8.5, 10318 BP, Karl Storz, Germany). (B) Coagulation suction tube (10390 BN, Karl Storz, Germany). (C) The cautery was performed on eight sites (including the membranous portion) of the tracheal segment for a length of 1 cm parallel to the trachea.

### Cuff overpressure intubation (COI) with an over-sized tube

A cuffed silicone endotracheal tube with an internal diameter of 9 mm (ID9)/outer diameter of 12 mm (OD12), and length of 33 cm (HS-ET-WC9.0W/NC9.0W, Hyupsung Medical, Korea) was selected as the over-sized tube, because tubes larger than ID9/OD12 were technically difficult to intubate. The proximal tip of the tube was placed 5 cm deeper than the upper incisor of the pig to ensure the cuff of the tube was located at least 5 cm below the vocal cords. A homemade pressure measuring device was used to monitor and maintain cuff overpressure (200– 500 mmHg) (Figure 2). The tracheal tube used in the present study was unable to withstand cuff pressures higher than 500 mmHg, limiting the maximum load of the cuff to 500 mmHg. Due to the laboratory operating time constraint, the cuff overpressure intubation period was limited to a maximum of 8 hours.

**Figure 1.**
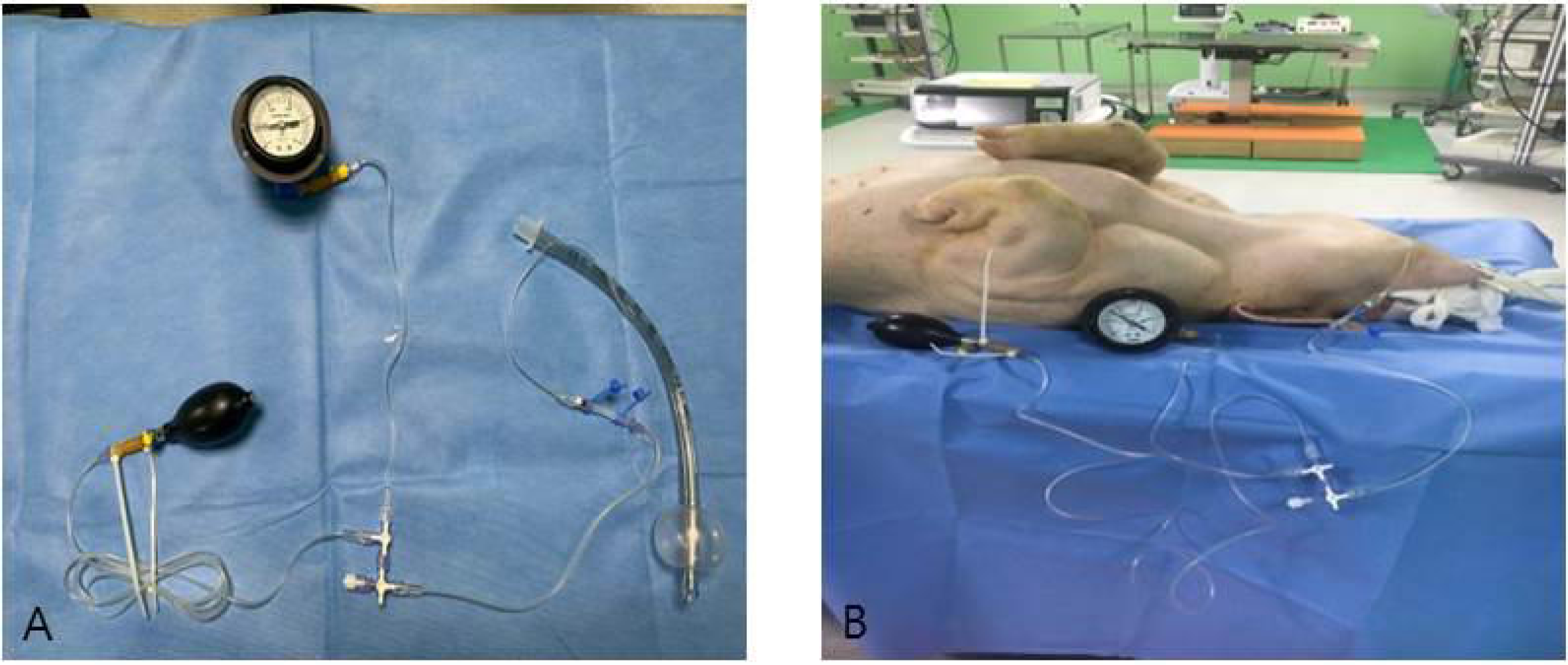
Homemade cuff-pressure measuring device. (A) The device is composed of a three-way connector, aneroid barometer, and air bulb. (B) After the cuff pressure was raised to the target pressure via an air bulb, general anesthesia was maintained. The pressure gauge was checked every 10 minutes and air was injected into the air bulb if it dropped below the target pressure.

### Tracheal stenosis

The tracheal lumen was examined using a flexible bronchoscope (MAF-TM, Olympus, Tokyo, Japan) on a weekly basis and the tracheal diameter was calculated considering the opening diameter of the biopsy forceps. The degree of tracheal stenosis was determined by the reduction of the tracheal lumen cross-section area: (S − s)/S × 100%, where “s” is the tracheal cross-sectional area after the intervention and “S” is the area before the intervention. A reduction of at least 50% of the measured airway cross-sectional area after intervention indicated a successful model. During the observation period, pigs exhibiting any signs of dyspnea, pain, costal retraction, or significant tracheal stenosis were euthanized using KCl injection. Pigs without any signs of breathing difficulty or fatal tracheal stenosis were euthanized on Day 21. After euthanasia, tracheas were removed for hematoxylin and eosin staining to observe any histological changes.

### Study groups

Experiments were performed on 14 pigs divided into three groups: tracheal cautery (TC) group (n = 3), cuff overpressure intubation (COI) group (n = 3), and COI-TC combination group (n = 8) (Table 1). In the TC group (n = 3), one pig (Pig TC1) received 60 watts electrocautery, and the other two pigs (Pig TC2 and TC3) received 40 watts electrocautery. In the COI group (n = 3), prolonged intubation with cuff overpressure was applied to the pigs after endotracheal tube (ID9/OD12) insertion. Pigs (Pig COI1, COI2, and COI3) were intubated for 8 hours with 200, 400, and 500 mmHg cuff pressure, respectively. In the COI-TC combination group (n = 8), immediately after COI (ID9/OD12 tracheal tube) at various cuff pressures (200–500 mmHg) and intubation durations (0.5–8 hours), electrocautery (40 watts) was performed on the tracheal mucosa where the cuff was located. In the TC group, if stenosis did not occur, repeated cautery was performed. However, no further intervention was allowed in the other groups.

**Table 1.** Details of the intubation duration, cuff pressure, tracheal cautery, and results of the tracheal stenosis model

| Pig No.<br>(n=14) | Tracheal inner diameter on day 0, mm | Tracheal cross-sectional area, mm<sup>2</sup> | Cuff overpressure intubation (COI) with ID 9.0 (OD 12) tube, day 0 |  | Tracheal cautery (TC), watt |  |  | Tracheal inner diameter, mm |  | Tracheal cross-sectional area, mm<sup>2</sup> |  | Stenosis, % |  | Length of stenosis, mm | Survival period, days |
| --- | --- | --- | --- | --- | --- | --- | --- | --- | --- | --- | --- | --- | --- | --- | --- |
|  |  |  | Cuff pressure, mmHg | Intubation duration, hour | Day 0 | Day 7 | Day 14 | Day 7 | Day 14 | Day 7 | Day 14 | Day 7 | Day 14 |  |  |
| TC1 | 16 | 201 | — | — | 60 | — | — | 13 | — | 133 | — | 34 | — | 15 | 7 |
| TC2 | 17 | 227 | — | — | 40 | 40 | — | 14 | 10 | 154 | 79 | 32 | 65 | 18 | 21 |
| TC3 | 18 | 254 | — | — | 40 | 40 | — | 15 | 11 | 177 | 95 | 31 | 63 | 20 | 21 |
| COI1 | 17 | 227 | 200 | 8 | — | — | — | 16 | 16 | 201 | 201 | 11 | 11 | 15 | 21 |
| COI2 | 16 | 201 | 400 | 8 | — | — | — | 15 | 15 | 177 | 177 | 12 | 12 | 14 | 21 |
| COI3 | 16 | 201 | 500 | 8 | — | — | — | 15 | 16 | 177 | 201 | 12 | 0 | 20 | 21 |
| COI-TC1 | 15 | 177 | 500 | 8 | 40 | — | — | 5 | — | 20 | — | 89 | — | 25 | 7 |
| COI-TC2 | 17 | 227 | 500 | 4 | 40 | — | — | 7 | 6 | 38 | 28 | 83 | 88 | 30 | 14 |
| COI-TC3 | 18 | 254 | 400 | 4 | 40 | — | — | 8 | 7 | 50 | 38 | 80 | 85 | 25 | 14 |
| COI-TC4 | 17 | 227 | 400 | 2 | 40 | — | — | 9 | 9 | 64 | 64 | 72 | 72 | 25 | 14 |
| COI-TC5 | 18 | 254 | 400 | 1 | 40 | — | — | 11 | 10 | 95 | 79 | 63 | 69 | 20 | 21 |
| COI-TC6 | 16 | 201 | 400 | 0.5 | 40 | — | — | 11 | 10 | 95 | 79 | 53 | 61 | 22 | 21 |
| COI-TC7 | 17 | 227 | 200 | 1 | 40 | — | — | 12 | 11 | 113 | 95 | 50 | 58 | 20 | 21 |
| COI-TC8 | 18 | 254 | 200 | 0.5 | 40 | — | — | 15 | 14 | 177 | 154 | 31 | 40 | 15 | 21 |

## RESULTS

The results of the 14 pigs in the three groups are summarized in Table 1. Detailed results of the groups are as follows.

### Tracheal cautery (TC) group

In the TC group (n = 3), the pig that received 60 watts cautery (Pig TC1) died suddenly 7 days after cautery. Autopsy showed that the pig had 34% tracheal stenosis. In the other two pigs that received 40 watts cautery (Pig TC2 and TC3), only 31–32 % stenosis had developed by post-cautery Day 7. Repeated cautery (40 watts) was applied to these two pigs on post-cautery Day 7. At 2 weeks after the initial cautery and 1 week after repeated cautery, hyperplasic granulation tissue grew in the tracheal wall and caused significant stenosis (63–65 %) in these pigs (Figure 3 and Table 1).

**Figure 3.**
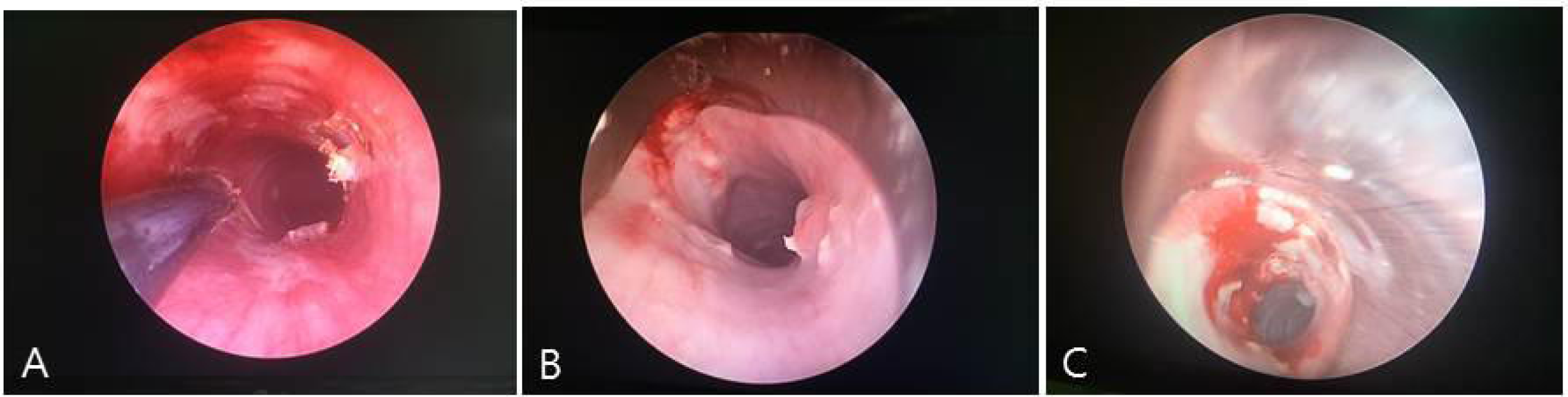
Tracheal cautery (TC) group. More than one cautery was needed to induce tracheal stenosis. (A) On Day 0, 40 watts cautery was performed on Pig TC2. (B) Day 7 in Pig TC2, stenosis was not significant (32 %), therefore, repeated cautery was performed. (C) Day 14 in Pig TC2, significant stenosis (65%) had developed as a result of granulation tissue hyperplasia.

### Cuff overpressure intubation (COI) group

In the COI group (Pig COI1, COI2, and COI3), 8-hour prolonged intubation was performed with cuff overpressures of 200, 400, and 500 mmHg, respectively. Immediately after prolonged intubation with cuff overpressure, the tracheal mucosa showed redness and sloughing. These ischemia-induced mucosal changes were increased with higher cuff pressures. However, no significant tracheal stenosis occurred during the 2 weeks after intervention (COI) (Figure 4 and Table 1).

**Figure 4.**
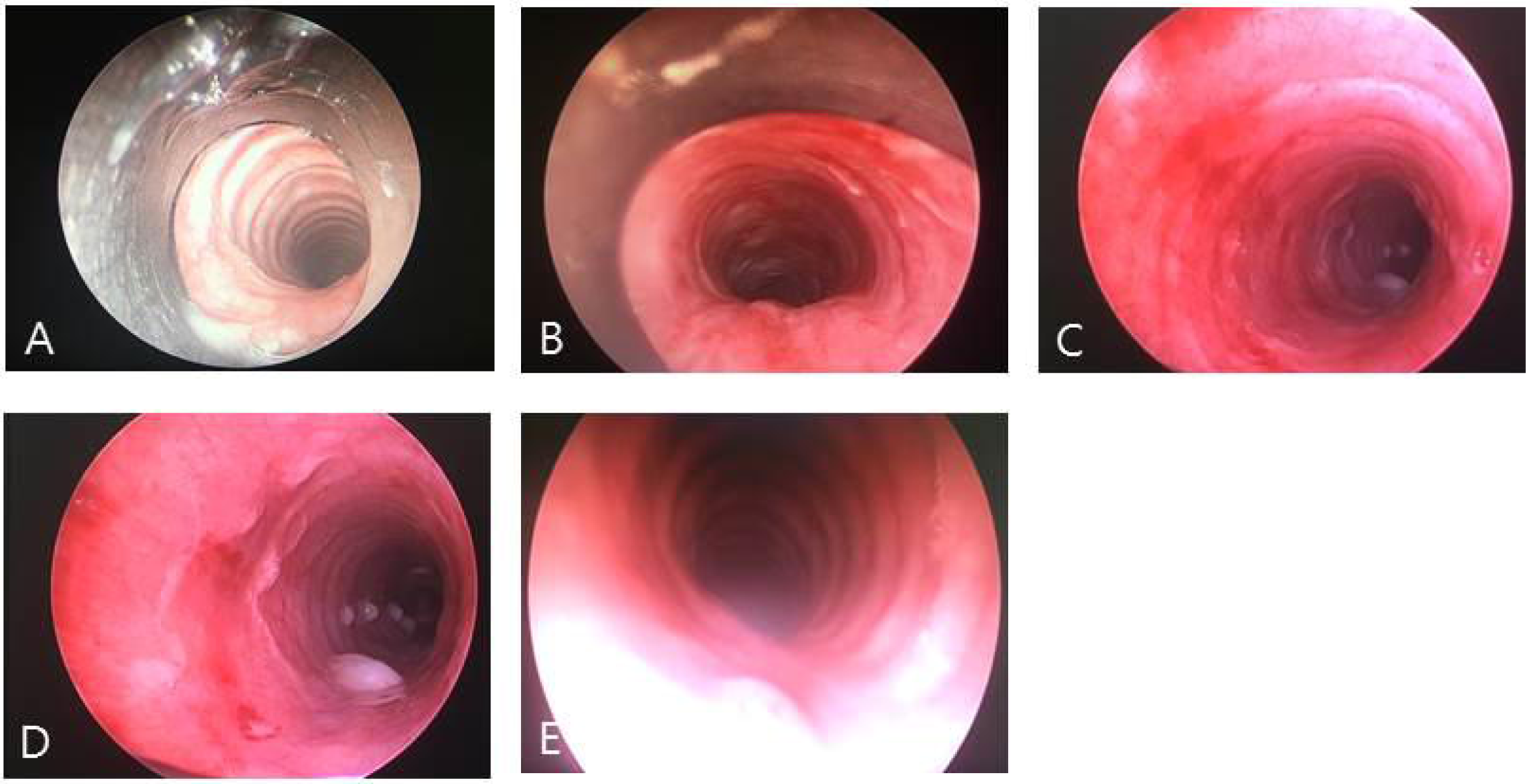
Cuff overpressure intubation (COI) group. The higher the cuff-pressure, the greater the mucosal change (redness and sloughing) that was noted as reflected by the ischemia-induced inflammatory reaction. However, these mucosal changes were not followed by tracheal stenosis. (A) Day 0 in Pig COI1, before COI. (B) Day 0, immediately after 8-hour 200 mmHg COI. (C) Day 0 in Pig COI2, immediately after 8-hour 400 mmHg COI. (D) Day 0 in Pig COI3, immediately after 8-hour 500 mmHg COI. (E) Day 7 in Pig COI3, significant stenosis had not developed (12%).

### COI-TC group

In the COI-TC group, eight pigs were divided into three subgroups according to the degree of cuff overpressure: two (Pig COI-TC1 and COI-TC2) at 500 mmHg, four (Pig COI-TC3–COI-TC6) at 400 mmHg, and two (Pig COI-TC7 and COI-TC8) at 200 mmHg. In the 500-mmHg subgroup, two pigs (Pig COI-TC1 and COI-TC2) underwent 8-hour and 4-hour prolonged intubation, respectively. In the 400-mmHg subgroup, four pigs (Pig COI-TC3–COI-TC6) underwent 4-hour, 2-hour, 1-hour, and 0.5-hour prolonged intubation, respectively. In the 200-mmHg subgroup, two pigs (Pig COI-TC7 and COI-TC8) underwent 1-hour and 0.5-hour prolonged intubation, respectively. After COI, 40 watts cautery was performed on the tracheal mucosa where the cuff was located.

In the 500-mmHg subgroup, one pig (Pig COI-TC1, 8-hour intubation) died suddenly 7 days after COI-TC. Autopsy showed that the pig had 89% tracheal stenosis. Another pig (Pig COI-TC2, 4-hour intubation) in this subgroup showed 83% stenosis and 88% stenosis at 7 and 14 days after COI-TC, respectively (Figure 5A). The pig was euthanized because of severe dyspnea accompanying costal retraction (Table 1).

**Figure 5.**
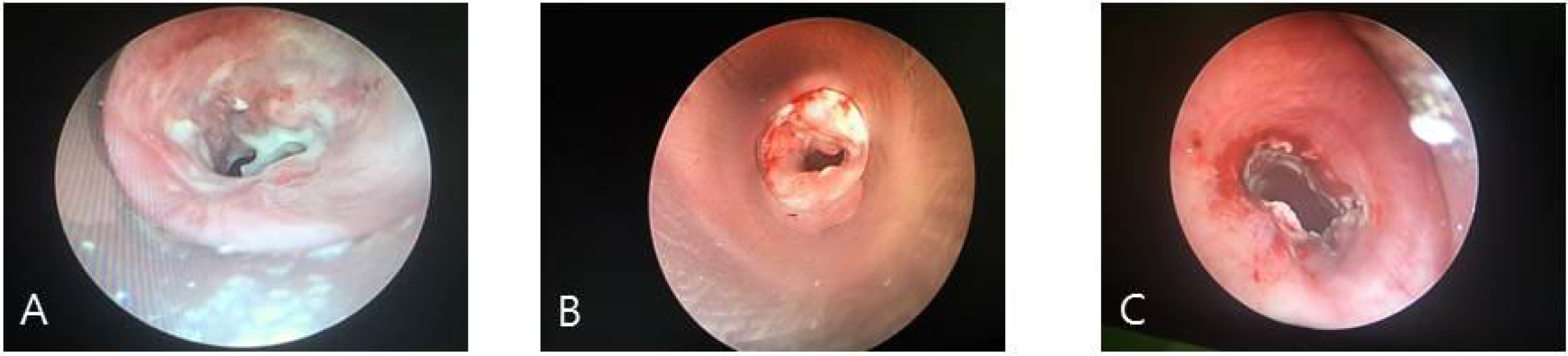
COI-TC group. The combination of prolonged COI and tracheal injury (tracheal cautery) allows rapid establishment of tracheal stenosis. Tracheal stenosis was increased in proportion to the cuff pressure and tracheal intubation time. (A) Day 7 in Pig COI-TC2, significant stenosis (83%) was noted. (B) Day 7 in Pig COI-TC3, significant stenosis (80%) was noted. (C) Day 7 in Pig COI-TC7, significant stenosis (50%) was noted.

In the 400-mmHg subgroup, all pigs (Pig COI-TC3–COI-TC6, 0.5–4-hour intubation) showed significant stenosis (53–80%) at 7 days after COI-TC (Figure 5B). The degree of stenosis was proportional to the tracheal intubation time. At 14 days after COI-TC, the stenoses were similar or slightly increased (61–85 %). Pig COI-TC3 and COI-TC4 were euthanized before 21 days because of severe dyspnea accompanying costal retraction (Table 1).

In the 200-mmHg subgroup, one pig (Pig COI-TC7, 1-hour intubation) showed 50% stenosis and 58% stenosis at 7 and 14 days after COI-TC, respectively (Figure 5C). The other pig (Pig COI-TC8, 0.5-hour intubation) in this subgroup showed no significant tracheal stenosis during the 2 weeks after intervention (Table 1).

Figure 6 shows the histologic findings from Pig COI-TC1 after 7 days of intervention. Severe hyperplasia of granulation tissue and inflammation of the tracheal wall, including the cartilage, were noted.

**Figure 6.**
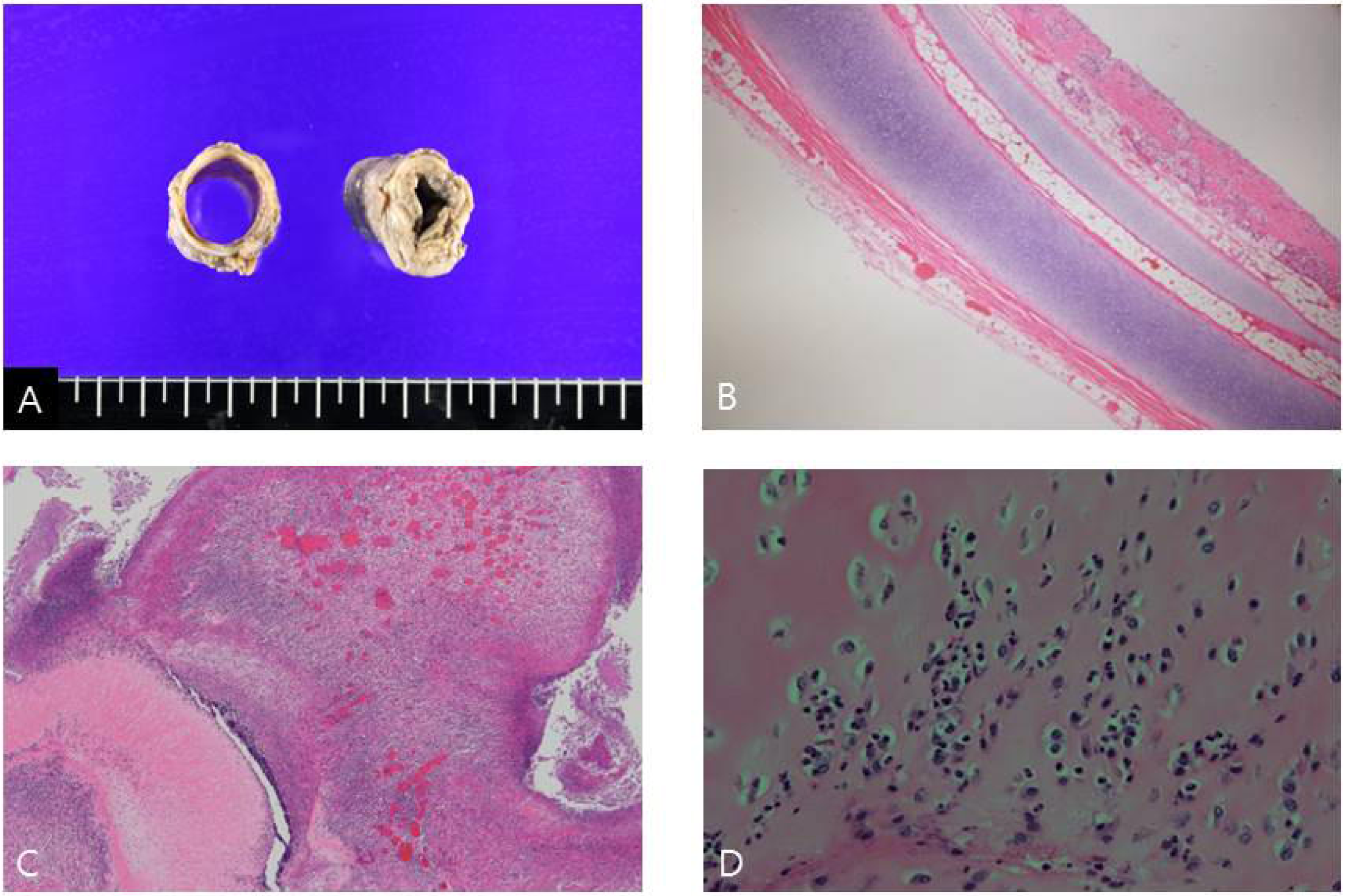
Histology of a representative case of COI-TC group (Day 7 in Pig COI-TC1). (A) Gross tracheal appearance of a normal segment (left) and stenotic segment (right). (B) Mucosal, submucosal, and cartilaginous structures in the normal tracheal segment (×40). (C) Inflammatory exudate and proliferative granulation tissue in the stenotic tracheal segment (×40). (D) Inflammatory cellular infiltration in the cartilage of the stenotic tracheal segment (×400).

## DISCUSSION

This animal study revealed that the combination of cuff overpressure intubation and tracheal injury (tracheal cautery) allows rapid establishment of tracheal stenosis in pigs. When the cauterization (40 watts) was performed immediately after over-sized intubation (ID9/OD12) for 1 hour with a cuff pressure of 200 mmHg or more, significant tracheal stenosis occurred in 7 days. In addition, the degree of tracheal stenosis tended to increase in proportion to cuff pressure and tracheal intubation time. This animal model is technically simple and reproducible, and the human tracheal stenosis mechanism is used. This will be of great help in developing new therapies for CAO.

CAO is a possibly life-threatening condition that can be caused by a number of malignant and non-malignant diseases (6). The best treatment is surgery; however, this is contraindicated in many cases; therefore a bronchoscopic treatment, such as an airway stent, is performed. Bronchoscopic treatment is not inherently a complete treatment but rather can lead to complications (e.g., for airway stents: migration, mucostasis, and granulation tissue formation). Therefore it is continuously improving and new treatments are being developed (3-6). The application of new treatments in humans requires preclinical studies with appropriate animal models. However, previous animal models (dog, pig, and rabbit) of tracheal stenosis take a relatively longer time (3–8 weeks) to develop and require complex processes, and the developmental mechanisms differ somewhat from the actual mechanisms of human tracheal stenosis (7-10). Human tracheal stenosis is caused by excessive cuff pressure, over-sized intubation, and tracheal damage (11, 13, 15-18). The present study was conducted to determine how fast tracheal stenosis can be produced by applying the above three mechanisms individually or simultaneously.

In our study, through simultaneous application of cuff overpressure intubation and tracheal cautery, successful airway stenosis was induced in seven out of eight pigs (in COI-TC group) within a week. Furthermore, the degree of tracheal stenosis increased in proportion with the cuff pressure and tracheal intubation time. Depending on the degree of cuff overpressure, the minimum required intubation time (immediately before cautery) was different (400 mmHg, 30 minutes; 200 mmHg, 1 hour). Cuff overpressure alone did not result in significant tracheal stenosis. It did not occur even when the tracheal intubation was maintained for 8 hours at a cuff pressure of 500 mmHg. Tracheal cautery induced airway stenosis, but repeated procedures and a relatively longer time (14 days) were required.

Previous studies have shown that if the cuff pressure is higher than the capillary pressure during intubation, submucosal blood flow stops causing ischemic damage to the tracheal wall. If the cuff pressure is maintained at 100 mmHg for 4 hours, the tracheal cartilage becomes damaged and inflamed (19, 20). Inflammation of the tracheal wall could induce airway narrowing during the healing process via granulation tissue and fibrotic tissue formation (11-13). Su et al. produced tracheal stenosis in dogs in a relatively shorter period of time (in 2 weeks) using prolonged intubation with cuff overpressure (14). Their model required prolonged intubation for at least 24 hours at a cuff pressure of 200 mmHg using an ID7.5 tracheal tube. They did not apply any other manipulation, such as cautery. In our study, 8-hour prolonged intubation with a much higher cuff pressure (500 mmHg) did not induce tracheal stenosis. Perhaps, prolonged intubation is more important than cuff overpressure per se in inducing tracheal wall inflammation. However, long-term general anesthesia involves high cost. Sometimes it may not be possible due to the constraints on laboratory operating time, such as in our animal laboratory. We had to find a way to induce tracheal stenosis and the artificial injury (cautery) was attempted in the tracheal mucosa where the cuff pressure was applied.

Electrocautery has been used to induce tracheal stenosis in animals alone or together with another intervention (10, 15). In this study, stenosis was established in 2–3 weeks. We tried both cautery alone and combined with cuff overpressure. The cautery power was set at 40–60 watts, which is a commonly used range in humans. At 60 watts, however, sudden death occurred with no stenosis induction; accordingly, subsequent experiments were conducted at 40 watts. In our study, to achieve significant tracheal stenosis using cautery alone, two or more sessions of cautery and at least 14 days were needed. In contrast, significant stenosis occurred successfully in 7 days in the cautery plus cuff overpressure combination group. Furthermore, by adjusting the cuff pressure and intubation duration, the degree of stenosis could be predicted. In our opinion, there is a synergistic effect on the induction of tracheal inflammation when both cautery and cuff overpressure intubation are performed simultaneously.

Animal models of airway stenosis are used for the investigation and development of new effective treatments for CAO. To achieve this, the establishment of animal models should not take too long and complex or expensive methods are also undesirable. In addition, the degree of induced stenosis can be predicted such that stenosis that is significant but not life-threatening is desirable. Finally, it would be better if it was induced using the same mechanisms leading to human tracheal stenosis. Our animal model takes only 7 days to establish (faster than all previous studies), is technically simple and cost effective (induction of tracheal stenosis even with short anesthesia time), and uses the same mechanisms as those underlying human benign tracheal stenosis. In addition, the degree of airway narrowing can be predicted; thus, researchers can induce tracheal stenosis as desired.

The histology of tracheal stenosis in our experiment (COI-TC group) showed acute inflammation of the entire airway wall (including the tracheal cartilage) and granulation tissue overgrowth. Previous studies also have similar histologic findings (18, 21, 22). Researchers have described that significant tracheal stenosis seems to occur when inflammation of the airway wall spreads to tracheal cartilage beyond the mucosa and submucosa. It is thought that, in our study, cuff overpressure alone did not induce sufficient damage to the tracheal cartilage.

Our study had some limitations. First, the number of pigs was small. To generalize our findings, replication studies may be necessary. Second, there was a lack of results for histological analysis. If the histological analysis was performed more closely according to the group and time point of pigs, we could better identify what is needed to establish tracheal stenosis.

## CONCLUSION

Through the present study, we found that the combination of cuff overpressure intubation and tracheal injury (tracheal cautery) allows rapid establishment of tracheal stenosis in pigs (within 7 days). In addition, the degree of tracheal stenosis increased in proportion to the cuff pressure and tracheal intubation time. Using this protocol, researchers can induce the desired degree of tracheal stenosis. This animal model is technically straightforward and uses mechanisms similar to those underlying human (benign) tracheal stenosis, making it a good experimental tool for developing new therapies for CAO.

## Acknowledgement

The study was funded by the National Research Foundation of Korea (NRF-2017R1C1B5076493).

## Ethics

The study protocol was approved by the Institutional Animal Care and Use Committee at Pusan National University Hospital (Approval Number: PNUYH-2017-043).

